# Spatial arrangement drastically changes the neural representation of multiple visual stimuli that compete in more than one feature domain

**DOI:** 10.1101/692541

**Authors:** Steven Wiesner, Ian W. Baumgart, Xin Huang

**Affiliations:** Department of Neuroscience, School of Medicine and Public Health, Physiology Graduate Training Program, McPherson Eye Research Institute, University of Wisconsin - Madison, WI 53705, U.S.A.

## Abstract

Natural scenes often contain multiple objects and surfaces. However, how neurons in the visual cortex represent multiple visual stimuli is not well understood. Previous studies have shown that, when multiple stimuli compete in one feature domain, the evoked neuronal response is biased toward the stimulus that has a stronger signal strength. Here we investigate how neurons in the middle temporal (MT) cortex of macaques represent multiple stimuli that compete in more than one feature domain. Visual stimuli were two random-dot patches moving in different directions. One stimulus had low luminance contrast and moved with high coherence, whereas the other had high contrast and moved with low coherence. We found that how MT neurons represent multiple stimuli depended on the spatial arrangement of the stimuli. When two stimuli were overlapping, MT responses were dominated by the stimulus component that had high contrast. When two stimuli were spatially separated within the receptive fields, the contrast dominance was abolished. We found the same results when using contrast to compete with motion speed. Our neural data and computer simulations using a V1-MT model suggest that the contrast dominance found with overlapping stimuli is due to normalization occurring at an input stage fed to MT, and MT neurons cannot overturn this bias based on their own feature selectivity. The interaction between spatially separated stimuli can largely be explained by normalization within MT. Our results revealed new rules on stimulus competition and highlighted the impact of hierarchical processing on representing multiple stimuli in the visual cortex.

**SIGNIFICANCE STATEMENT:** Previous studies have shown that the neural representation of multiple visual stimuli can be accounted for by a divisive normalization model. By using multiple stimuli that compete in more than one feature domain, we found that luminance contrast has a dominant effect in determining competition between multiple stimuli when they were overlapping but not spatially separated. Our results revealed that neuronal responses to multiple stimuli in a given cortical area cannot be simply predicted by the population neural responses elicited in that area by the individual stimulus components. To understand the neural representation of multiple stimuli, rather than considering response normalization only within the area of interest, one must consider the computations including normalization occurring along the hierarchical visual pathway.

## Introduction

In natural scenes, multiple visual stimuli are often present in a local spatial region. While it is generally well understood how neurons in the visual cortex encode a single stimulus, how neurons encode multiple visual stimuli within their receptive fields (RFs) remains to be elucidated. Because visual perception depends critically on the integration and segregation of multiple visual stimuli (Braddick, 1993), understanding the neural representation of multiple stimuli is of significant importance.

The middle temporal (MT) cortex is an extrastriate brain area that is important for visual motion processing (Britten, 2003; Born and Bradley, 2005; Park and Tadin, 2018). Neurons in area MT receive feedforward inputs from direction-selective neurons in V1 (Movshon and Newsome, 1996) and have RFs about ten times larger in size than those of V1 neurons at the same eccentricities (Gattass and Gross, 1981; Albright and Desimone, 1987). Previous studies have shown that neuronal responses in area MT elicited by multiple moving stimuli follow a sub-linear summation of the responses elicited by the individual stimulus components (Snowden et al., 1991; Qian and Andersen, 1994; Recanzone et al., 1997; Ferera and Lisberger, 1997; Britten and Heuer, 1999; Heuer and Britten, 2002; McDonald et al., 2014), consistent with a model of divisive normalization (Simoncelli and Heeger, 1998; Britten and Heuer, 1999; Carandini and Heeger, 2011).

Work in our laboratory has shown that the direction tuning curves of MT neurons to overlapping random-dot stimuli moving transparently in different directions can also be described as a weighted sum of the responses elicited by the individual stimulus components (Xiao et al., 2014; Xiao and Huang, 2015). When two stimulus components have different signal strengths in one feature domain, defined either by motion coherence or luminance contrast, MT neurons pool the stimulus component that has a stronger signal strength with greater weight (Xiao et al., 2014). The response bias in MT toward the stimulus component that has a stronger signal strength can be accounted for by a descriptive model of divisive normalization (Xiao et al., 2014), similar to the contrast normalization model used to describe neuronal responses in V1 (Carandini et al., 1997; Busse et al. 2009).

However, natural scenes contain multiple visual stimuli that often differ in more than one feature domain. For example, one stimulus may have a stronger signal strength in feature A but a weaker signal strength in feature B, whereas another stimulus may have a weaker signal strength in feature A but a stronger signal strength in feature B. In this case, it is unclear which stimulus has an overall stronger signal strength and, more generally, how visual stimuli with multiple competing features interact within neurons’ RFs.

One possibility is that, to neurons in a given brain area, the overall signal strength of a visual stimulus is reflected in the evoked responses of a population of neurons in that area. Due to divisive normalization within that area, a neuron may weigh a visual stimulus more strongly if the population neural response elicited by that stimulus is greater than the population response elicited by a competing stimulus. Alternatively, how neurons in a given brain area weigh multiple competing stimuli may be the result of neural computations occurring in multiple stages along the hierarchical visual pathway and may not be explained by simply considering the population neural responses elicited by the individual stimulus components in the area of interest.

Here, we investigate the rule by with neurons in area MT encode multiple moving stimuli that compete in more than one feature domain. We found that MT responses to multiple stimuli changed drastically when the spatial arrangement of the visual stimuli was varied. Our results revealed how visual stimuli that differ in multiple feature domains interact within neurons’ RFs and shed light on how the neuronal responses in a given cortical area are shaped by neural processing along the hierarchical visual pathway.

## Materials and Methods

Two male adult rhesus monkeys (*Macaca mulatta*) were used in the neurophysiological experiments. Experimental protocols were approved by the Institutional Animal Care and Use Committee of UW-Madison and conform to U.S. Department of Agriculture regulations and to the National Institutes of Health guidelines for the care and use of laboratory animals. Procedures for surgical preparation and electrophysiological recordings were routine and similar to those described previously (Xiao et al., 2015). A head post and a recording cylinder were implanted during sterile surgery with the animal under isoflurane anesthesia. For electrophysiological recordings from neurons in area MT, we took a vertical approach and used tungsten electrodes (1-3 MΩ, FHC). We identified area MT by its characteristically large portion of directionally selective neurons, small RFs relative to those of neighboring medial superior temporal cortex (area MST), its location at the posterior bank of the superior temporal sulcus, and visual topography of the RFs (Gattass and Gross, 1981). Electrical signals were amplified and single units were identified with a real-time template-matching system and an offline spike sorter (Plexon). Eye position was monitored using a video-based eye tracker (EyeLink, SR Research) with a rate of 1000 Hz.

### Visual stimuli and experimental procedure

Stimulus presentation and data acquisition were controlled by a real-time data acquisition program “Maestro” (https://sites.google.com/a/srscicomp.com/maestro/home). Visual stimuli were presented on a 25-inch CRT monitor at a viewing distance of 63 cm. Monitor resolution was 1024 × 768 pixels, with a refresh rate of 100 Hz. Stimuli were generated by a Linux workstation using an OpenGL application that communicated with an experimental control computer. The luminance of the video monitor was measured with a photometer (LS-110, Minolta) and was gamma-corrected.

Visual stimuli were achromatic random-dot patches presented within a circular aperture with a diameter of 3°. Individual dots were squares of 2 pixels extending 0.08° on each side, and each random-dot patch had a dot density of 2.7 dots/deg^2^. The dots had a luminance of either 79 or 22 cd/m^2^, presented on a uniform background with a luminance of 10 cd/m^2^, which gives rise to a Michelson contrast of either 77.5% or 37.5%. Random dots in each patch moved within the stationary aperture in a specified direction. The motion coherence of each random-dot patch was set to either 100% or 60%. To generate a random-dot patch moving at N% of motion coherence (after Newsome and Pare 1988; Britten et al. 1992), N% of the “signal” dots were selected to move coherently, while the rest of the dots referred to as the “noise” dots were repositioned randomly within the aperture. Random selections of the “signal” and “noise” dots occurred at each monitor frame. Therefore, a given dot would switch back and forth between a signal dot and a noise dot. The lifetime of each dot was as long as the motion duration.

In each experimental trial, the monkey maintained fixation within a 1° × 1° electronic window around a small fixation point. After a neuron was isolated, we first characterized its direction selectivity by interleaving trials of a 30° × 27° random-dot patch, moving in different directions at a step of 45° and at a speed of 10°/s. The direction selectivity and preferred direction (PD) were determined on-line using MATLAB (MathWorks). We then characterized the speed tuning of the neuron using a random-dot patch moving at different speeds (1, 2, 4, 8, 16, 32, or 64°/s) in the neuron’s PD. Using a cubic spline, the preferred speed (PS) of the neuron was taken as the speed that evoked the highest firing rate in the fitted speed tuning curve. Next, we used a series of 5° × 5° random-dot patches moving in the PD and at the PS of the neuron to map the neuron’s RF. The location of the patch was randomized and the screen was tiled in 5° steps. The RF map was interpolated at 0.5° intervals, and the location giving rise to the highest firing rate was taken as the center of the RF.

In the main experiments, the visual stimuli appeared after the monkey maintained fixation for 200 ms. To separate the neuronal responses to the stimulus motion from those due to the stimulus onset, the visual stimuli were first turned on and remained stationary for 200 ms before they started to move for 500 ms. The visual stimuli were then turned off. The monkeys maintained fixation for an additional 200 ms after the stimulus offset. In some stimulus trials, two random-dot patches that moved in different directions, referred to as two stimulus components, were presented simultaneously. The direction separation between two stimulus components was fixed at 90°. We varied the vector averaged (VA) direction of the bi-directional stimulus around 360° to characterize the response tuning curve. The two stimulus components were either overlapping in one of two locations (site a or b) within the RF, or they were spatially separated within the RF, one centered at site a and the other at site b, with at least 1° gap between the borders of the two random-dot patches (illustrated in Fig. 1). In other trials, only one stimulus component was presented at either site a or site b and the direction was varied to characterize the tuning curve to the stimulus component. For the majority of the experiments, the VA and component directions were varied in a step of 15°. In a small set of experiments, the directions were varied in a step of 30°. The trials presenting bi-directional stimuli and individual stimulus components were randomly interleaved.

**Figure 1.**
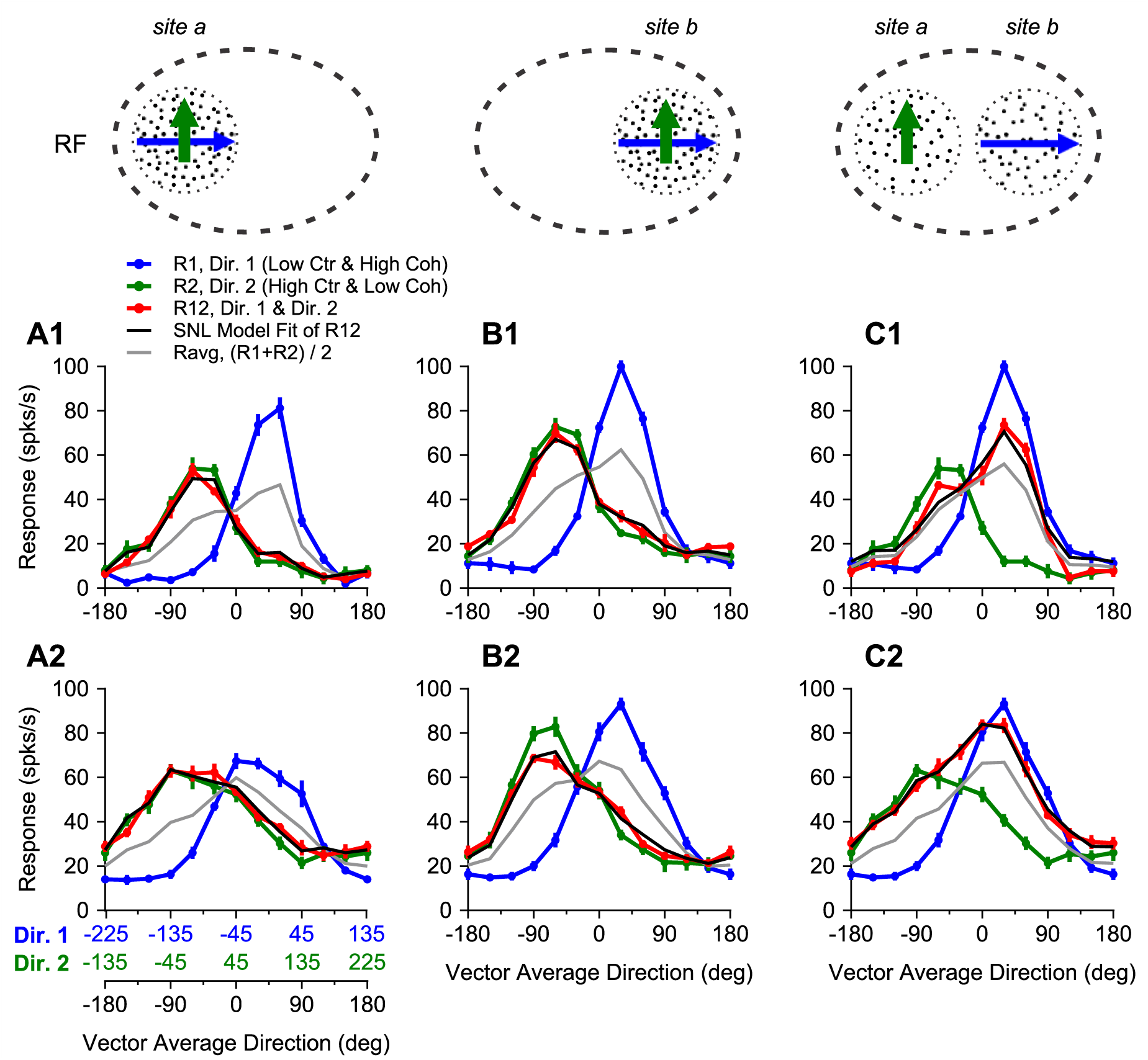
The response tuning curves of two example MT neurons to overlapping (A, B) and spatially-separated stimuli (C). Visual stimuli were achromatic random-dot patches moving in two directions separated by 90°. The “low contrast & high coherence” component (shown in blue arrow) moved at the clockwise side of the two component directions, whereas the “high contrast & low coherence” component (shown in green arrow) moved in the direction at the counter-clockwise side. The X-axis labeled in black indicates the vector average direction of the bi-directional stimuli. The X-axes labeled in blue and green (**A2**) indicate the direction of the “low contrast & high coherence” component (Dir. 1) and the direction of “high contrast & low coherence” component (Dir. 2), respectively. The three X-axes are aligned such that the component directions shown in blue and green correspond to the directions of the two stimulus components at each vector average direction. **A1-C1:** Response tuning curves from one neuron. **A2-C2:** Response tuning curves from another neuron. The responses elicited by the bi-directional stimuli are shown in red (R12). The SNL model fits of R12 are shown in black. Error bars represent standard errors.

In the first experiment, one random-dot patch, referred to as the “low contrast & high coherence” component, had a luminance contrast of 37.5% and a motion coherence of 100%. The other random-dot patch, referred to as the “high contrast & low coherence” component, had a luminance contrast of 77.5% and a motion coherence of 60%. Both stimulus components moved at the same speed, which was set at the neuron’s PS if it was below 10°/s, or at 10°/s if the PS was at or greater than 10°/s. Note that when a random-dot patch moved at 60% coherence in a given direction, the visual stimulus was different from a situation where 60% of the dots always moved coherently and the rest of the 40% of dots always moved randomly. Because the random selection of signal and noise dots occurred at each monitor frame in our stimuli, a noise dot at one frame may turn into a signal dot in the next frame and move in the coherent direction. Perceptually, it is difficult to segregate the noise dots from the signal dots of the same stimulus component. The noise dots of the “high contrast & low coherence component” are not an independent entity and do not appear to interfere with the coherence of the “low contrast & high coherence” component perceptually.

In the second experiment, we set the motion coherence of both random-dot patches to 100% but used different speeds for the two stimulus components. One random-dot patch, referred to as the “low contrast & faster speed” component, had a luminance contrast of 37.5% and moved at 10°/s. The other random-dot patch, referred to as the “high contrast & slower speed” component, had a luminance contrast of 77.5% and moved at 2.5°/s.

### Data analysis

Response firing rate was calculated during the period of 500-ms stimulus motion and averaged across repeated trials. We fitted the raw direction tuning curves for the bi-directional stimuli and the individual stimulus components using splines at a resolution of 1°. We then rotated the spline-fitted tuning curve to the bi-directional stimuli so that the VA direction of 0° was aligned with the PD of each neuron. In the first experiment, the responses of each neuron to the bi-directional stimuli and individual stimulus components were normalized by the maximum response to the “low contrast & high coherence” component. In the second experiment, the responses of each neuron were normalized by the maximum response to the faster speed component. We averaged the rotated and normalized tuning curves across neurons to obtain population-averaged tuning curves.

To quantify the relationship between the responses elicited by the bi-directional stimuli and those elicited by the individual stimulus components, we fitted the direction tuning curves using a summation plus nonlinear interaction (SNL) model (Eq. 1), which has been shown to provide a better fit of MT responses elicited by bi-directional stimuli than a linear weighted summation model (Xiao et al., 2014).

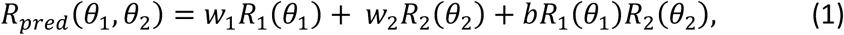

where *R_pred_* is the response to the bi-directional stimuli predicted by the model; *θ_1_* and *θ_2_* are the two component directions; *R_1_* and *R_2_* are the measured component responses elicited by the two stimulus components when presented alone; *w_1_* and *w_2_* are the response weights for *R_1_* and *R_2_*, respectively; and *b* is the coefficient of multiplicative interaction between the component responses. To determine whether the response elicited by the bi-directional stimuli showed a significant bias toward one of the two stimulus components, we compared the response weights *w_1_* and *w_2_* using either a paired t-test or a Wilcoxon signed-rank test.

We also fitted the response tuning curves to the bi-directional stimuli using a few variants of a divisive normalization model (Carandini and Heeger, 2011) (see Results). The model fits were obtained using the constrained minimization tool ‘fmincon’ (MATLAB) to minimize the sum of squared error.

To evaluate the goodness of fit of a model for the response tuning curve to the bi-directional stimuli, we calculated the percentage of variance (PV) accounted for by the model as:

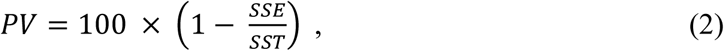

where SSE is the sum of squared errors between the model fit and the neuronal data, and SST is the sum of squared differences between the data and the mean of the data (Morgan et al., 2008).

### V1-MT Model

We adapted a computational model proposed by Simoncelli and Heeger (1998) (http://www.cns.nyu.edu/~lcv/MTmodel/) to reconstruct our visual stimuli and to simulate the neuronal response tuning to the bi-directional stimuli that were either overlapping or spatially separated. The model contained several consecutive stages, which can be interpreted as V1 simple, V1 complex, and MT (Simoncelli and Heeger, 1996; Rust et al., 2006). Based on the dimensions of video monitor and viewing distance in our neurophysiological experiments, 1° of visual angle corresponds to 21 pixels. The random-dot patch in our model simulations had a circular aperture with a diameter of 63 pixels (i.e. 3°) and the same dot density as used in our experiments. Each dot had a size of 2 × 2 pixels.

We set the RFs of model neurons by Gaussian convolutional filters (Table 1). We estimated the size of the RF for each neuron type by summing the lengths of the incorporated filters. For the spatially-separated stimuli, we set a blank gap between the two stimulus components as the RF size of the V1 complex neuron, which is 1.2°, to ensure that no V1 neuron would be driven by both stimulus components. We generated direction-selective neuron populations that approximately tiled a sphere in the frequency domain. We tuned the contrast response functions by adjusting *C_50_* values for V1 and MT neurons. These *C_50_* values were represented in the model as σ^2^ in the normalization equation (Eq. 3), which was applied to both V1 complex cell and MT stages of the model (adapted from Simoncelli and Heeger 1998 and Rust et al., 2006).

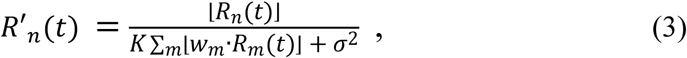

where *R_n_(t)* represents the *nth* neuron’s linear filter response; *R’_n_(t)* represents the normalized response of either V1 complex cell or MT neuron; ⌊ ⌋ denotes half-wave rectification; *K* represents the strength of normalization, which was set as 1-*σ^2^*; *m* represents the *nth* neuron’s normalization pool; *w* represents the Gaussian spatial weighting profile of the normalization pool, with a standard deviation of *SD_norm_*. The model parameters for V1 and MT stages are defined in Table 1. We fitted the model contrast response functions to neural data from V1 and MT as described in Sclar et al. (1990). Similarly, we tuned coherence responses by varying the spatial scale of the normalization pool (*m*), the weighting profile within the pool (*w*), and the size of the V1 linear RF. The MT coherence response function was fitted to data replotted from Figure 1C in Britten and Newsome (1998). We are not aware of published neural data on V1 coherence response function. So the parameters for V1 model neurons were varied to simulate our MT responses to bi-directional stimuli without a constraint on V1 coherence response function. The same model parameters were used for the overlapping and spatially separated conditions.

**Table 1.** Model parameters for V1 and MT neurons

| Model parameters | V1 stage | MT stage |
| --- | --- | --- |
| RF size (pixels) | 15 (simple cell)<br>25 (complex cell) | 211 |
| Standard Deviation (SD) of Gaussian RF profile (pixels) | 0.73/2.19/6.57 (3 scales for simple cells)<br>7 (complex cell) | 53 |
| $\sigma^2 (C_{50})$ | 0.0016 | 0.000049 |
| Size of normalization pool $m$ (pixels) | 38 | 153 |
| $SD_{norm}$ (pixels) | 5 | 52 |
| Baseline activity | 0 | 0.1 |

We explored several variants of the model architecture. The model parameters were fitted after each architectural manipulation. The following changes enabled the model to better capture the trends of the stimulus competition found in our neural data. First, we used area-normalized Gaussian functions to set the weights for the spatial pooling and local population normalization. Second, multiple frequency scales for V1 simple cells were computed by tripling the standard deviation of the underlying 3^rd^ order derivative Gaussian, similar to the doubling suggested in Simoncelli and Heeger (1998) – this change was made after spectral analysis of stimuli showed that a wider range of scales was necessary to capture motion at lower coherence. Third, V1 afferent weights were not adjusted to zero mean, allowing MT neurons to have variable proportions of positive and negative inputs. Finally and importantly, rectification and static nonlinearity were applied to the MT stage after spatial pooling and before normalization, which is physiologically plausible and provides a better fit of our neural data.

## Results

We asked the question of how neurons in extrastriate area MT represent multiple visual stimuli that compete in more than one feature domain. To address this question, we conducted neurophysiological experiments and computer simulations. We recorded from isolated single neurons in area MT of two macaque monkeys while they performed a fixation task. Visual stimuli were two random-dot patches moving simultaneously in different directions within the RFs. In the first experiment, we used luminance contrast and motion coherence as two competing features. One stimulus had high contrast but moved with low coherence, whereas the other stimulus had low contrast but moved with high coherence (see Methods). We manipulated the spatial arrangement of the visual stimuli to investigate the contributions of earlier visual areas and area MT in mediating the competition between multiple stimuli. In a second experiment, we used luminance contrast and motion speed as two competing features. We first present the results from the neurophysiological experiments and then computer simulations.

### Neurophysiological experiments

We measured the direction tuning curves of MT neurons in response to two stimuli that had competing visual features and moved simultaneously in different directions. Our dataset includes recordings from 76 MT neurons, 43 from monkey G and 33 from monkey B. We set the angular separation between the motion directions of two individual stimuli, referred to as the stimulus components, at 90° and varied the VA direction of the stimuli. In the first experiment, one stimulus component had a low contrast of 37.5% and moved at a high motion coherence of 100%. The other component had a high contrast of 77.5% and moved at a low coherence of 60%. Figure 1 shows the direction tuning curves of two representative neurons. The red curve shows the neuronal response elicited when both stimulus components were present, as a function of the VA direction of the two stimulus components. The green and blue curves show the neuronal responses elicited by the individual stimulus components when presented alone. The tuning curves of the component responses are arranged such that, at each VA direction, the data points on the green and blue curves correspond to the responses elicited by the individual stimulus components of that VA direction (note the color-coded abscissas for the component directions in Fig. 1A2).

For the two example neurons, the peak response of the direction tuning curve to the “low contrast & high coherence” component alone (shown in blue) was greater than that to the “high contrast & low coherence” component (shown in green) (Fig. 1). This is expected since MT neurons are sensitive to motion coherence within a large coherence range (Britten et al., 1993), whereas their contrast response function saturates at a low luminance contrast (Sclar et al., 1990). Consequently, the average of the response tuning curves to the two stimulus components (shown in gray) was biased toward the “low contrast & high coherence” component. Surprisingly, we found that when the two stimulus components were overlapping, the neuronal responses elicited by the bi-directional stimuli were strongly biased toward the “high contrast & low coherence” component (Fig. 1A). This response bias was robust and occurred when we placed the overlapping stimuli at a different site within the RF (Fig. 1B).

Two overlapping visual stimuli could stimulate not only the RFs of single MT neurons but also the RFs of single V1 neurons. The response bias toward the “high contrast & low coherence” component may be caused by the neural processes within area MT, or alternatively inherited from earlier visual areas such as V1. To determine the contribution of earlier visual areas to the response bias, we placed two stimulus components at different locations within the RF of a given MT neuron. The two stimulus components were separated by a gap of at least 1° (illustrated in Fig. 1C). With this spatial arrangement, the RF of a single V1 neuron could only be stimulated by one of the two stimulus components, whereas the RF of an MT neuron could still be stimulated by both components. We found that the response tuning to the bi-directional stimuli changed drastically when stimulus components were spatially separated. MT responses elicited by the bi-directional stimuli no longer showed a bias toward the “high contrast & low coherence” component, but roughly followed a scaled average of the component responses (Fig. 1C).

Figure 2 shows the tuning curves averaged across 70 MT neurons. The population-averaged response elicited by the “low contrast & high coherence” component moving in the PD of each neuron, aligned to 0°, was significantly greater than that elicited by the “high contrast & low coherence” component moving in the PD (one-tailed paired t-test, p = 4.1×10^−7^). However, when the two stimuli were overlapping, the population response elicited by the bi-directional stimuli was almost completely biased toward the weaker “high contrast & low coherence” component, regardless of the spatial location within the RF (Fig. 2A and 2B). The bias toward the “high contrast & low coherence” component at a given VA direction was in a manner of “higher-contrast-take-all”. For example, at a VA direction of 45° where the “low contrast & high coherence” component moved in the PD (0°) and the “high contrast & low coherence” component moved in 90° (indicated by a dotted line in Fig. 2A), the bi-directional response closely followed the much weaker response elicited by the “high contrast & low coherence” component. When the two stimulus components were spatially separated within the RF, the strong bias toward the “high contrast & low coherence” component was abolished (Fig. 2C). The population response to the bi-directional stimuli now showed roughly equal weighting of the responses elicited by the individual stimulus components.

**Figure 2.**
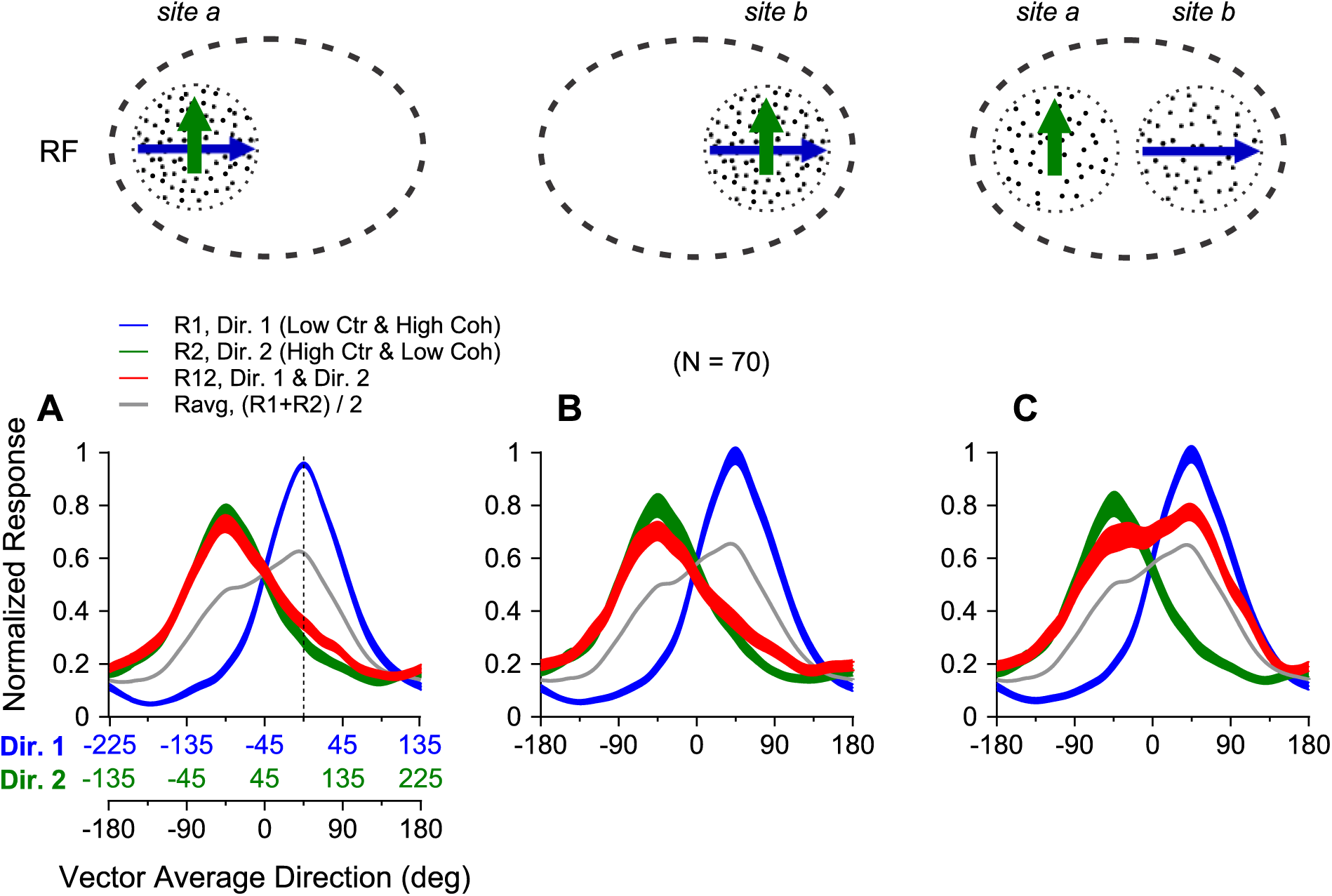
Population-averaged tuning curves to the bi-directional stimuli (red) and the unidirectional stimulus components (blue and green). The vector average direction of the bi-directional stimuli and the directions of individual stimulus components are labeled in the corresponding X-axes (**A**), following the same convention as in Figure 1. The direction of 0° was aligned with each neuron’s PD before the tuning curves were averaged across neurons. The stimulus components were overlapping at site *a* (**A**) or site *b* (**B**), or spatially separated (**C**) within the RFs. The width of each tuning curve represents the standard error. The average of the responses to the two stimulus components is shown in gray.

The SNL model (see Eq. 1 in Methods) provided an excellent fit of the MT responses elicited by the bi-directional stimuli, illustrated by the black curves in Figure 1. Across our neuron population the model fit accounted for, on average, 83% of the response variance (see Methods). Figure 3 compares the response weights for the two stimulus components obtained from the SNL model fits. In the overlapping condition, the mean response weight *w_2_* for the “high contrast & low coherence” component was significantly greater than the weight *w_1_* for the “low contrast & high coherence” component (one-tailed paired t-test, p = 1.9×10^−45^ for *site a*, p = 2.5×10^−28^ for *site b*) (Fig. 3A). Nearly all data points, each representing the result from one neuron, were below the unity line. The mean response weight for the “high contrast & low coherence” component was 0.97 (std = 0.24), whereas the mean weight for the “low contrast & high coherence” component was 0.23 (std = 0.25), indicating a dominant effect of the “high contrast & low coherence” component in determining the neuronal response to the bi-directional stimuli.

**Figure 3.**
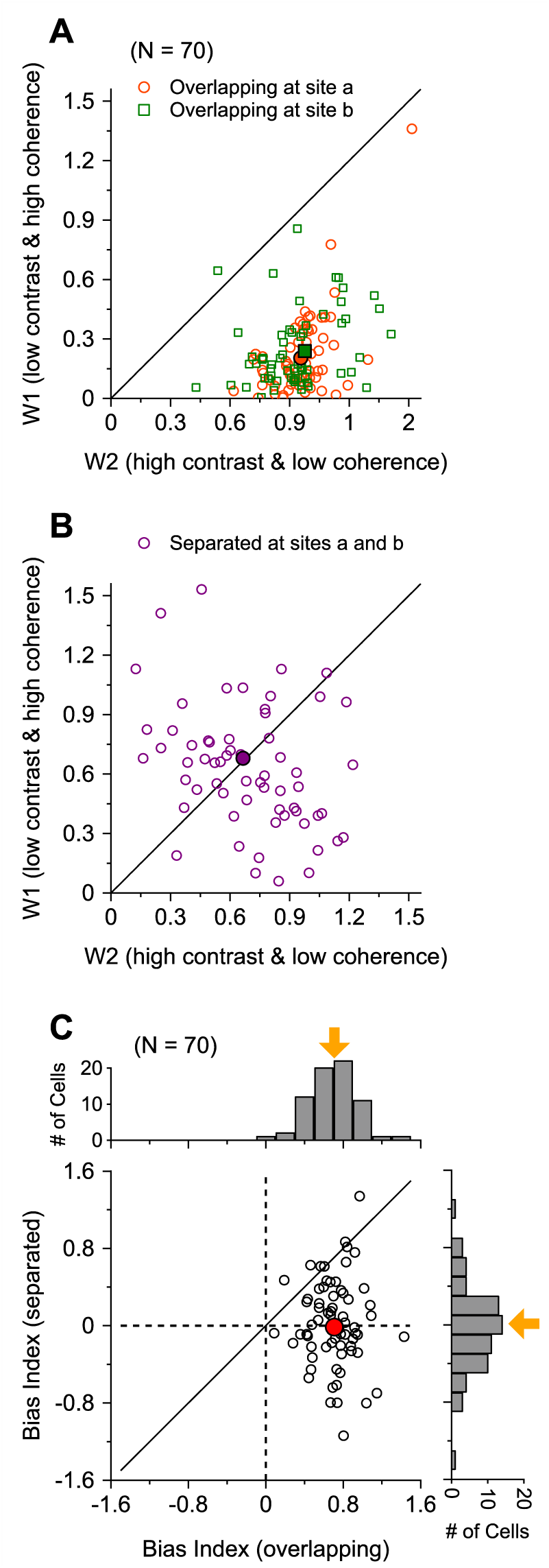
The effect of the spatial arrangement of the bi-directional stimuli on the response weights for the stimulus components. Each dot represents the result from one neuron. Comparing the response weights for the “low contrast & high coherence” component (ordinate) with the “high contrast & low coherence” component (abscissa) under the overlapping (**A**) and the spatially separated (**B**) conditions. **C.** Comparing the bias indices between the spatially separated (ordinate) and overlapping (abscissa) conditions. The histograms in C show the distributions of the bias index for the overlapping (top) and spatially separated (right) conditions.

When the two stimulus components were spatially separated within the RF, the response weights changed significantly, becoming symmetrically distributed relative to the unity line (Fig. 3B). The spread of weights in the spatially-separated condition is larger than that in the overlapping condition. The mean weight for the “high contrast & low coherence” component decreased to 0.66 (std = 0.32), whereas the mean weight for the “low contrast & high coherence” component increased to 0.68 (std = 0.43). The mean weights for the two components were no longer different (paired t-test, p = 0.8) but were significantly greater than 0.5 of response averaging (t-test, p < 0.001).

To quantify the response bias toward an individual stimulus component, we calculated a bias index (BI):

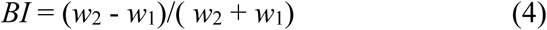

A positive value of the index indicates a bias toward the “high contrast & low coherence” component. Figure 3C shows how this bias index changes with the spatial arrangement of the visual stimuli. In the overlapping condition, the mean BI is 0.73 (std = 0.23), which is significantly greater than 0 (one-tailed t-test, p = 7.5×10^−35^). In the spatially-separated condition, the mean BI is −0.01 (std = 0.95), which is not significantly different from 0 (p = 0.7). The mean BI obtained in the overlapping condition is significantly greater than that in the spatially-separated condition (one-tailed paired t-test, p = 4.7×10^−9^), indicating a change of the response bias when the spatial arrangement of the visual stimuli is altered.

We previously found that the tuning curves of some MT neurons to overlapping bi-directional stimuli can show a directional “side-bias” toward one of the two direction components (Xiao and Huang, 2015). A subgroup of neurons prefers the stimulus component at the clockwise side of two motion directions, whereas another group prefers the component direction at the counter-clockwise side. These response biases can occur even when both stimulus components have the same contrast and coherence. In the experiment shown in Figures 1-3, the “high contrast & low coherence” component always moved at the counter-clockwise side direction (Fig. 2A, 2B). Could the strong bias toward the “high contrast & low coherence” component in the overlapping condition be due to a biased neuron sample that happened to have a strong bias toward the direction component at the counter-clockwise side? To address this concern, we arranged the direction components differently.

Figure 4A and B show the averaged direction tuning curves of 15 MT neurons when the direction of the “high contrast & low coherence” component was placed at the counter-clockwise side under the overlapping and spatially separated conditions, as in Figure 2. When the “high contrast & low coherence” component was placed at the clockwise side of the two component directions, the responses of the same 15 neurons to the bi-directional stimuli still showed a strong bias toward the “high contrast & low coherence” component under the overlapping condition (Fig. 4C), and showed roughly equal weighting of the two components under the spatially-separated condition (Fig. 4D). Placing the “high contrast & low coherence” component at the clockwise or counter-clockwise side of the two component directions had no effect on the response bias, as measured by the bias index under the overlapping and spatially-separated conditions (Wilcoxon rank-sum test, p = 0.6).

**Figure 4.**
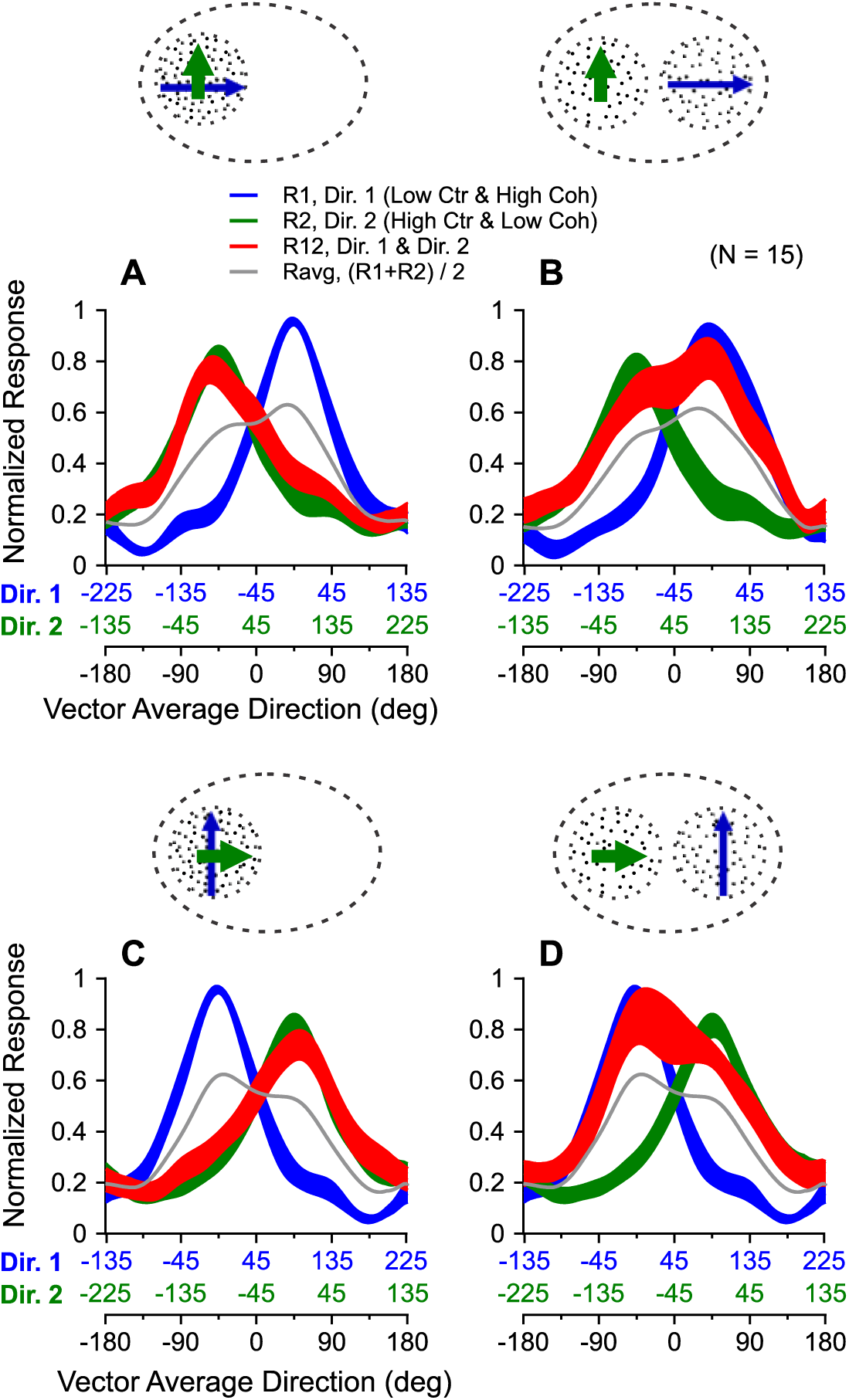
Control for the directional arrangement of the two stimulus components. **A, B.** Response tuning curves averaged across 15 MT neurons to the bi-directional stimuli and the stimulus components when the direction of the “high contrast & low coherence” component was placed at the counter-clockwise side of the two component directions, as in Figures 1 and 2. **C, D.** Response tuning curves averaged across the same 15 neurons when the direction of the “high contrast & low coherence” component was placed at the clockwise side of the two component directions. **A, C.** Overlapping condition. **B, D.** Spatially separated condition. Notice the switch of the values in the X-axes of the component directions, shown in blue and green, between A and C, as well as between B and D.

To shed light on the neural mechanisms underlying the response bias, we examined the timecourse of the neuronal responses in the overlapping and spatially separated conditions. Figure 5 shows the PSTHs calculated using a 10-ms time bin when either the “high contrast & low coherence” component or the “low contrast & high coherence” component moved in the PD. When stimuli were overlapping, as soon as MT neurons started to respond to the onset of the static stimuli (see Methods), the response elicited by both stimulus components already closely followed the “high contrast & low coherence” component, even before the onset of the stimulus motion (Fig. 5A, B). After the onset of the motion response, the neuronal response to the bi-directional stimuli continued to follow the response elicited by the “high contrast & low coherence” component throughout the motion period, regardless of whether the component moved in the PD and elicited a strong response (Fig. 5A), or 90° away from the PD and elicited a weak response (Fig. 5B). Since the strong bias towards the “high contrast & low coherence” component in the overlapping condition occurred at the very beginning of the stimulus onset, it is unlikely that the bias was due to selective attention (see Discussion).

**Figure 5.**
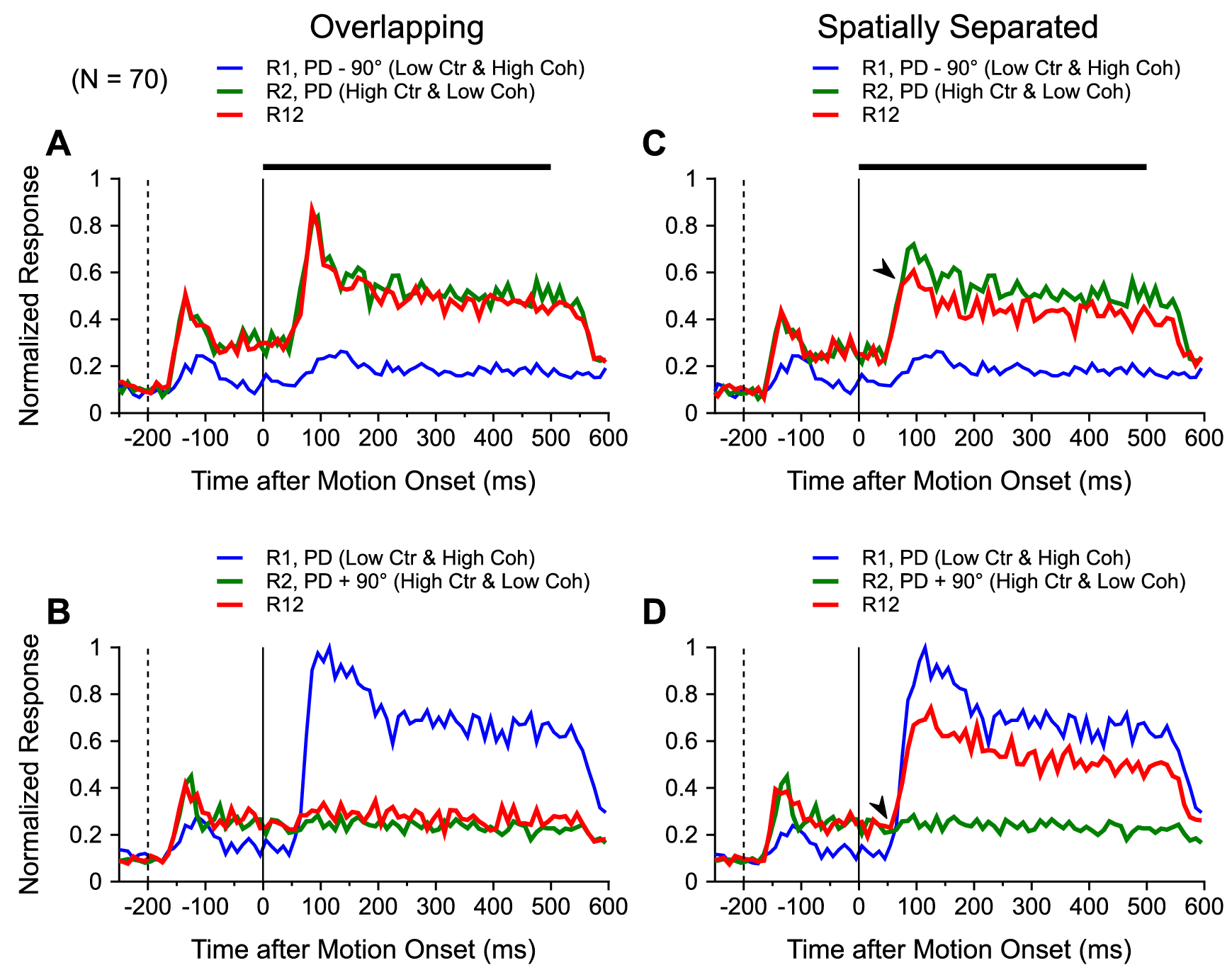
Timecourse of the neuronal responses to the bi-directional stimuli and the stimulus components. Peristimulus time histograms (PSTHs) were calculated using a 10-ms time bin and averaged across 70 neurons. **A, B.** The two stimulus components overlapped (at site *a*) within the RF. **C, D.** The two stimulus components were spatially separated within the RF. The dashed vertical lines at −200 ms indicate the onset of the static stimuli. The solid vertical lines at time 0 indicate motion onset. The solid horizontal bars shown in A and C indicate the stimulus motion period. The “high contrast & low coherence” component moved in the PD in A and C, and moved in a non-PD in B, D.

When stimuli were spatially separated, MT neurons also followed the “high contrast & low coherence” component in response to the onset of the static stimuli (Fig. 5C, D). After the motion onset, when the “high contrast & low coherence” component moved in the PD, the motion response elicited by the bi-directional stimuli initially followed the “high contrast & low coherence” component for ∼30 ms, and was then “pulled down” by the non-PD component (see the arrow in Fig. 5C). When the “high contrast & low coherence” component moved in the non-PD, the motion response elicited by the bi-directional stimuli followed the “high contrast & low coherence” component for ∼10 ms after the onset of the motion response to the PD component, and was then “pulled up” by the PD component (see the arrow in Fig. 5D). These results suggest that response normalization under the spatially separated condition takes 10∼30 ms to occur.

When two stimulus components overlap, the random dots from each component constitute only half of the total number of dots of the two moving surfaces. Could the strong response bias toward the “high contrast & low coherence” component be due to a reduction of the motion coherence of the “low contrast & high coherence” component when the stimuli overlapped? We think this is an unlikely explanation because overlapping reduces the percentage of the signal dots relative to the total number of dots for both stimulus components. In addition, our stimuli moved in two directions separated by 90°. Human observers can reliably segregate the two stimulus components at this angle separation and the “low contrast & high coherence” component still appears to move coherently. Overlapping does not change the relative coherence levels nor the perceived coherence of the two stimulus components. When overlapping random-dot stimuli have the same luminance contrast but move at different motion coherences, macaque MT response to both stimulus components is biased toward the high coherence component (Xiao et al., 2014), indicating that stimulus overlapping does not prevent the response bias toward the high coherence component given equal contrast.

To determine whether the dominance by the high-contrast component on MT responses elicited by overlapping stimuli occurs only when luminance contrast and motion coherence compete with each other, we conducted a second experiment using visual stimuli that differ in luminance contrast and motion speed. We previously found that when two overlapping random-dot patches moved in the same direction at different speeds, within a range of low to intermediate speeds, the responses of MT neurons elicited by the bi-speed stimuli was biased toward the faster speed component (X. Huang et al., unpublished data). Motivated by this finding, we used motion speed to compete with luminance contrast. As in the main experiment, the visual stimuli contained two random-dot patches moving in two directions separated by 90° and we varied the VA direction to measure the direction tuning curves. One stimulus component had a high luminance contrast of 77.5% and moved at a slower speed of 2.5°/s. The other stimulus component had a low luminance contrast of 37.5% and moved at a faster speed of 10°/s. Both stimulus components moved at 100% coherence and were either overlapping or spatially-separated within the RF of a given MT neuron as in the first experiment. We also measured the direction tuning curves when the two stimulus components both had high luminance contrast (77.5%) and moved at 2.5°/s and 10°/s, respectively, at 100% coherence.

We recorded from 13 MT neurons using these visual stimuli. Figure 6 shows the population-averaged tuning curves. When both stimulus components had high contrast, the peak response elicited by the faster (10°/s) stimulus component moving in the PD (i.e. 0°) was greater than that elicited by the slower (2.5°/s) component moving in the PD. The component responses are shown in green and purple in Figure 6A. When the two stimulus components were overlapping, the tuning curve elicited by both stimulus components (shown in red) is biased toward the faster stimulus component, more than what is predicted by the average of the component responses (shown in gray) (Fig. 6A). We fitted the direction tuning curves using the SNL model for each neuron (Eq. 1). The median response weight obtained by the model fit for the faster stimulus component (0.88) was significantly greater than the median weight (0.41) for the slower component (Wilcoxon signed-rank test, p = 7.3 × 10^−4^). This result extended our previous finding of the response bias toward the faster stimulus component for stimuli moving in the same direction (unpublished results) to stimuli moving in different directions.

**Figure 6.**
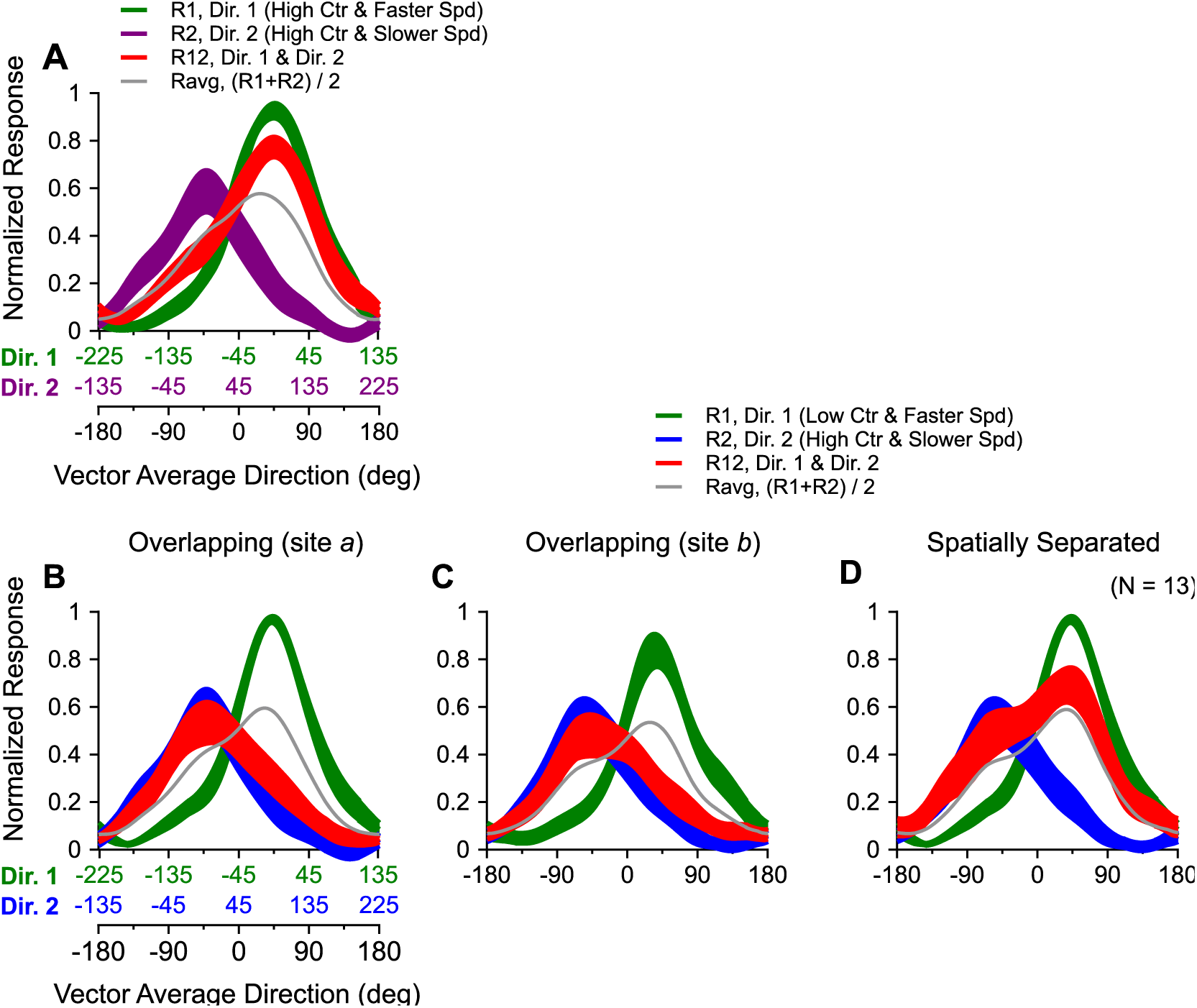
Averaged response tuning curves to two stimulus components that moved in different directions and at different speeds. Both stimulus components moved at 100% coherence. The response tuning to both stimulus components presented simultaneously is shown in red. The width of each tuning curve represents the standard error. The average of the component responses elicited by the individual stimulus components is shown in gray. **A.** Both stimulus components had high luminance contrast and were overlapping. **B-D.** The two stimulus components competed in luminance contrast and motion speed. The faster speed component had low luminance contrast, whereas the slower speed component had high luminance contrast. The stimulus components were overlapping at site *a* (**B**) or site *b* (**C**), or were spatially separated (**D**) within the RFs. The vector average direction of the bi-directional stimuli and the directions of individual stimulus components are labeled in the corresponding X-axes (**A, B**), following the same convention as in Figure 2.

When the overlapping stimuli moving at different speeds had different luminance contrasts, the responses elicited by both stimulus components showed a strong bias toward the “high contrast & slower speed” component, even though the peak response to this component alone was significantly weaker than that to the “low contrast & faster speed” component (Fig. 6B). We found the same result when the two stimulus components overlapped at a different site within the RF (Fig. 6C). Under the overlapping condition, the median response weight for the “high contrast & slower speed” component was 0.81, which was significantly greater than the median weight for the “low contrast & faster speed” component (0.17) (Wilcoxon signed-rank test, p = 2.4 × 10^−4^). Separating the two stimulus components spatially within the RF abolished the bias toward the “high contrast & slower speed” component (Fig. 6D). As the spatial arrangement of the stimulus components changed from overlapping to spatially separated, the median bias index (Eq. 4) decreased significantly from 0.65 to −0.08 (Wilcoxon signed-rank test, p = 0.0012). These results confirmed that luminance contrast has a dominant effect on MT responses elicited by overlapping stimuli, which is not unique to the competition between contrast and motion coherence. The spatial arrangement of visual stimuli can substantially change the competition between multiple stimuli within the RF.

### Fitting response tuning curve using the normalization model

Previous studies have shown that neuronal responses elicited by multiple stimuli in many brain areas can be described by a divisive normalization model (Carandini and Heeger, 2011). We asked whether our results could also be accounted for by response normalization. We first fitted the data using the following equation:

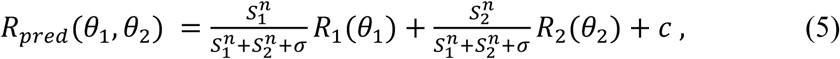

where *R_1_* and *R_2_* are the evoked direction tuning curves to the two stimulus components 1 and 2, respectively. *θ_1_* and *θ_2_* are the component directions. *S_1_* and *S_2_* represent the signal strengths of the “low contrast & high coherence” component and the “high contrast & low coherence” component, respectively. *R_pred_* is the model-predicted response elicited by both stimulus components presented simultaneously. *n, σ*, and *c* are model parameters with the constraints of *n* ≥ 1 and *c* > 0. Equations of the similar form have been used previously to describe normalization involving contrast, in which case the signal strength is simply the luminance contrast (Carandini et al., 1997; Busse et al., 2009; Xiao et al., 2014; Bao and Tsao, 2018). Since our visual stimuli competed in more than one feature domain, it was not obvious which stimulus component had an overall stronger signal strength. Because the brain has to make an inference of the signal strength based on the elicited neural responses, we assumed that the signal strength of a stimulus component, in the “eye” of MT neurons, is reflected in the neural responses elicited by that stimulus component moving in a fixed direction summed across a population of MT neurons that have different PDs evenly spanning 360°. This summed population response is invariant to the direction of the stimulus component, which is suitable for representing signal strength. Equivalently, the summed population neural response in MT can be approximated by summing the responses of each neuron elicited by stimulus component *i* moving in different directions spanning 360° and averaged across neurons in our data sample, e.g. to sum the population-averaged component responses across directions in Figure 2. This was how we calculated *Si,* (*i* =1, 2).

This normalization model (Eq. 5) failed to capture the response tuning to overlapping bi-directional stimuli, accounting for only 33% of the response variance (34% for site *a*, 32% for site *b*). The model performed better when stimuli were separated, accounting for 66% of the variance. We found similar results when using this model to fit the data from our second experiment, in which luminance contrast competed with motion speed. The model accounted for an average of 44% of the response variance (38% for site *a*, 50% for site b) when stimuli were overlapping, and 77% of the variance when stimuli were separated (Table 2).

**Table 2.** Fitting the direction tuning curves using the normalization model

| Visual Stimuli | $S_1$ | $S_2$ | Percentage of Variance Accounted for<br>(mean ± std) | | |
| --- | --- | --- | --- | --- | --- |
| Contrast vs. Coherence<br>(N = 70) | Low contrast & high coherence | High contrast & low coherence | Normalization<br>(Eq. 5) | Tuned Normalization<br>(Eq. 6) | Normalization with weighted Numerators<br>(Eq. 7) |
| Overlapping (site a) | 122.3 | 126.0 | 34 ± 18 | 34 ± 18 | 86 ± 16 |
| Overlapping (site b) | 128.3 | 130.9 | 32 ± 19 | 32 ± 19 | 81 ± 19 |
| Spatially Separated | 130.4 | 130.3 | 66 ± 24 | 68 ± 25 | 83 ± 17 |
| Contrast vs. Speed<br>(N = 13) | Low contrast & faster speed | High contrast & slower speed |  |  |  |
| Overlapping (site a) | 128.1 | 83.6 | 38 ± 21 | 39 ± 21 | 88 ± 14 |
| Overlapping (site b) | 113.3 | 81.3 | 49 ± 20 | 49 ± 20 | 84 ± 15 |
| Spatially Separated | 128.1 | 81.3 | 77 ± 20 | 77 ± 20 | 90 ± 5 |

It has been suggested that response normalization can be tuned, such that individual stimulus components contribute differently to normalization (Ni et al., 2012; Rust et al., 2006; also see Carandini et al., 1997). We therefore fitted our data using a tuned normalization equation:

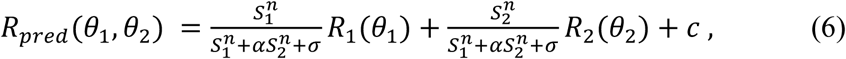

where α is a positive parameter that scales the contribution of *S_2_* with respect to *S_1_* to normalization. We found that introducing tuned normalization did not improve the model performance at all when stimuli were overlapping, accounting for an average of 33% of the response variance (34% for site *a*, 32% for site *b*). When stimuli were separated, the tuned normalization model accounted for 68% of the variance. We found the same results when fitting the data collected when contrast competed with speed (Table 2).

The poor fit of the responses under the overlapping condition by the standard normalization model (Eq. 5) can be understood because MT neurons showed a very strong bias toward the high contrast component, whereas *S_1_* and *S_2_* were similar. The tuned normalization was not able to improve the fit because, although it changed the relative contributions of the stimulus components to the normalization pool in the denominator, it kept the numerators in Equation 6 unchanged. Hence the relative weights for the two stimulus components did not change. To capture the strong bias toward the high contrast component in the overlapping condition, a weighting parameter is needed in the numerator. Accordingly, we fitted our results using the following equation:

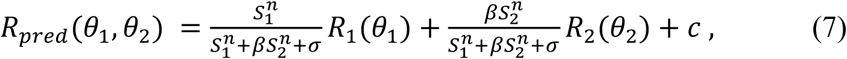

where *β* is a positive parameter and appears in both the numerator and the denominator. This parameter allows the relative response weights for the two stimulus components to vary. When *β* is greater than one, the response weight for the high contrast component (*R_2_*) is greater than that for the low contrast component (*R_1_*). As expected, this equation fitted the data well, accounting for >80% of the response variance for both the overlapping and spatially separated conditions (Table 2). However, the normalization model itself does not provide an explanation for why the response weight is greater for the high contrast component in the overlapping condition but not in the spatially separated condition.

### Computer simulations using a V1-MT model

Our spatially separated visual stimuli fall inside the RFs of single MT neurons, whereas only one of the stimulus components would fall inside the RFs of single V1 neurons. Hence, our spatially-separated visual stimuli can interact within the RFs of MT neurons but not V1 neurons. In contrast, the overlapping stimuli can interact within the RFs of both MT and V1 neurons. To explore the neural mechanisms underlying our physiological findings, we conducted computer simulations using a hierarchical feedforward model adapted from Simoncelli and Heeger (1998). This model consists of two processing stages corresponding to areas V1 and MT. Each stage carries out a series of computations including spatiotemporal filtering, spatial pooling, rectification, and divisive normalization. At the V1 stage, simple cells receive input directly from the visual stimulus and complex cells pool inputs from rectified and divisively normalized responses of V1 simple cells. At the MT stage, MT neurons pool inputs from V1 complex cells, followed by rectification and divisive normalization (Simoncelli and Heeger, 1998; Rust et al., 2006).

We generated random-dot visual stimuli that are similar to those used in our physiological experiments and simulated the neuronal responses in areas MT and V1. The visual stimuli and a simplified architecture of the model are illustrated in Figure 7. The diameter of each random-dot patch was 3°, extending 63 pixels. The RF sizes of model V1 and MT neurons, set by the sizes of the convolution filters, were 1.2° and 10° in diameter, respectively (see Methods). The populations of model neurons in V1 and MT stages approximately tiled a sphere in the spatiotemporal frequency domain, as in Simoncelli and Heeger’s model (1998). The RFs of V1 and MT neuron populations covered a region of the visual field that was 17.3° × 17.3°. In the overlapping condition, the apertures of two random-dot patches overlapped within the RFs (Fig. 7A). In the spatially-separated condition, the two random-dot patches were placed side by side, separated by a blank gap that was 1.2° wide, within the RFs of single MT neurons (Fig. 7B). In the overlapping condition, the V1 neurons whose RFs covered *site a* were activated by both stimulus components (Fig. 7A). In the spatially-separated condition, V1 neurons were activated by only one stimulus component, either at site *a* or site *b* (Fig. 7B).

**Figure 7.**
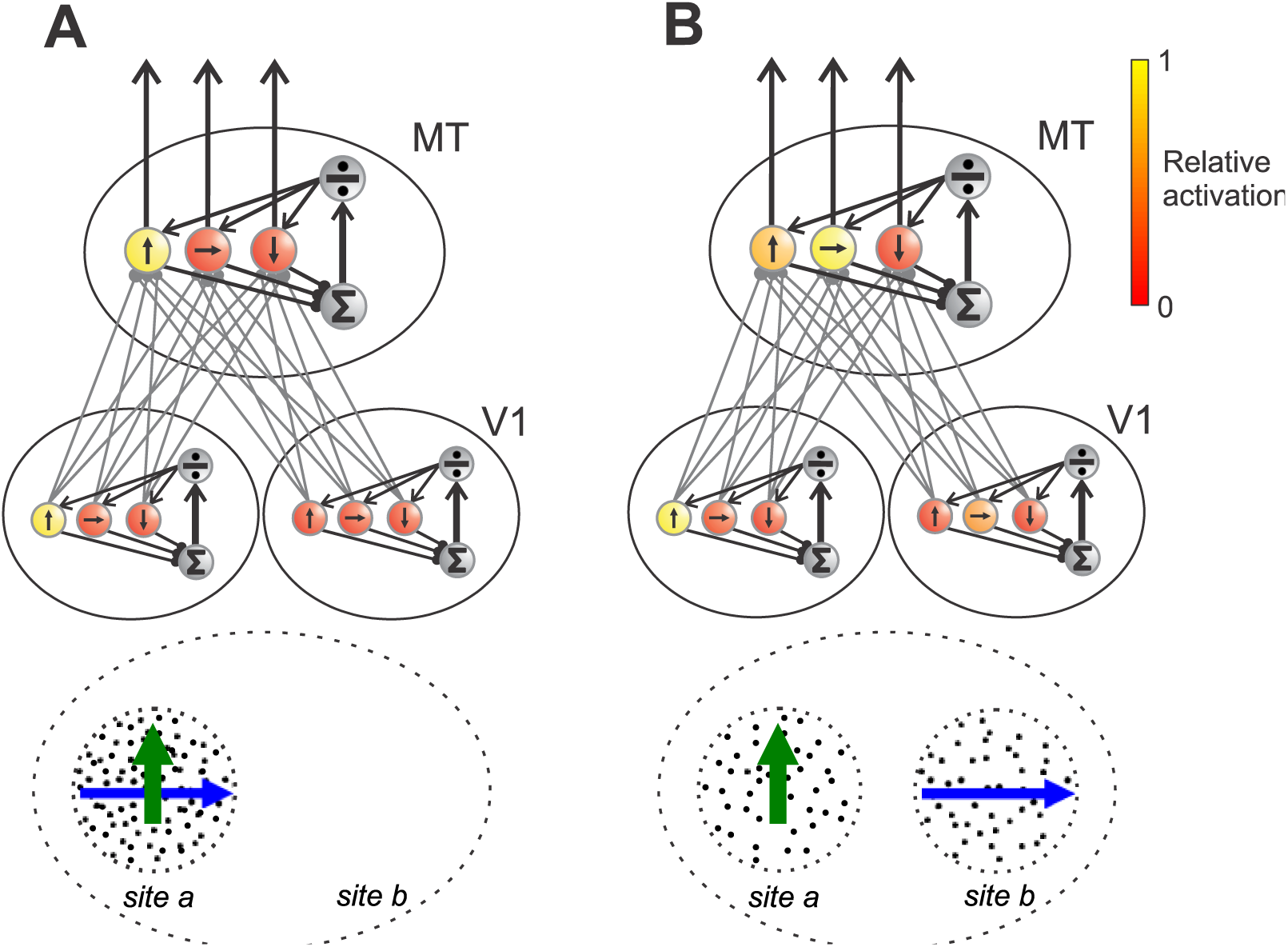
Illustration of a simplified architecture for the V1-MT model. Each MT neuron receives feedforward inputs from multiple neurons at the V1 stage. Responses are divisively normalized by the sum of local population activity at both V1 and MT stages. Each small circle represents a neuron and the black arrow inside the circle indicates the PD. The color of each circle indicates the response magnitude of the neuron. Yellow means maximum response and red means minimum response. Visual stimuli are illustrated below neural circuit as the input to the V1 stage. The green and blue arrows represent the “high contrast & low coherence” component and the “low contrast & high coherence” component, respectively. Two pools of neurons at the V1 stage that respond only to site *a* or site *b* respectively are illustrated. The RFs of the MT neurons are illustrated by the dotted ellipse and cover both site *a* and site *b*. **A.** Overlapping condition. **B.** Spatially separated condition.

We tuned the model parameters (see Methods) to match the experimentally measured contrast response functions of V1 and MT neurons (Sclar et al., 1990) and the coherence response function of MT neurons (Britten and Newsome, 1998). The simulated contrast response functions of V1 and MT neurons fitted the experimental data almost perfectly, and the simulated coherence response function of MT neurons also matched the data well (Fig. 8A-C). As far as we know, an experimentally measured coherence response function of V1 neurons has not been described previously. Our simulations show that V1 responses increased monotonically with the coherence level of moving random-dot stimuli (Fig. 8D). The model V1 neurons had lightly higher firing rates in response to low coherence stimuli and more trial-to-trial variability in comparison with the model MT neurons (Fig. 8C and D).

**Figure 8.**
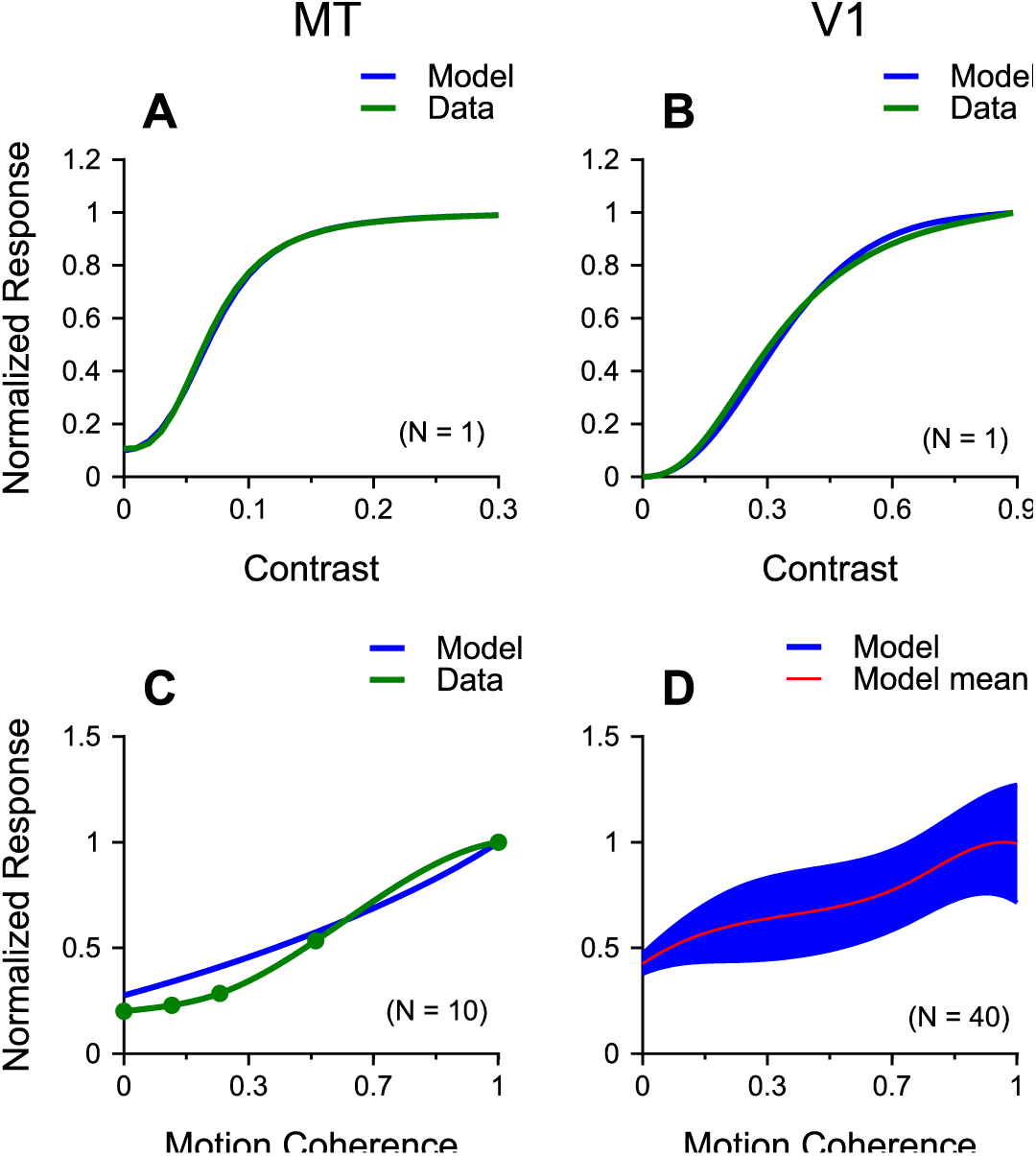
Contrast and coherence response functions of model V1 (**B, D**) and MT (**A, C**) neurons. **A, B.** Fitted contrast response functions to sinusoidal gratings for model neurons. Green curves are experimental data replotted from Sclar et al. (1990). **C**. Fitted coherence response function to high contrast random-dots for model MT neurons. Green dots are experimental data replotted from Britten and Newsome (1998). The green curve is the spline fit of the experimental data points. ***D***: Coherence response to high contrast random-dots for model V1 complex cells. The widths of the blue curves in C and D represent the standard deviation. N indicates the number of repeats for simulations. The stimulus dots were regenerated randomly for each simulation in C and D.

The MT responses elicited by our visual stimuli that competed between luminance contrast and motion coherence were well captured by the model. Consistent with our experimental data (Fig. 2), the tuning curve of model MT neurons to the “low contrast & high coherence” component had a greater peak response than that of the “high contrast & low coherence” component (Fig. 9A, B). In the overlapping condition, the simulated MT response elicited by the bi-directional stimuli was nearly completely biased toward the weaker “high contrast & low coherence” component (Fig. 9A), as found in the neural data. The model also captured the change of MT response tuning when visual stimuli were rearranged spatially. In the spatially-separated condition, the tuning curve of model MT neurons elicited by the bi-directional stimuli was no longer dominated by the “high contrast & low coherence” component (Fig. 9B).

**Figure 9.**
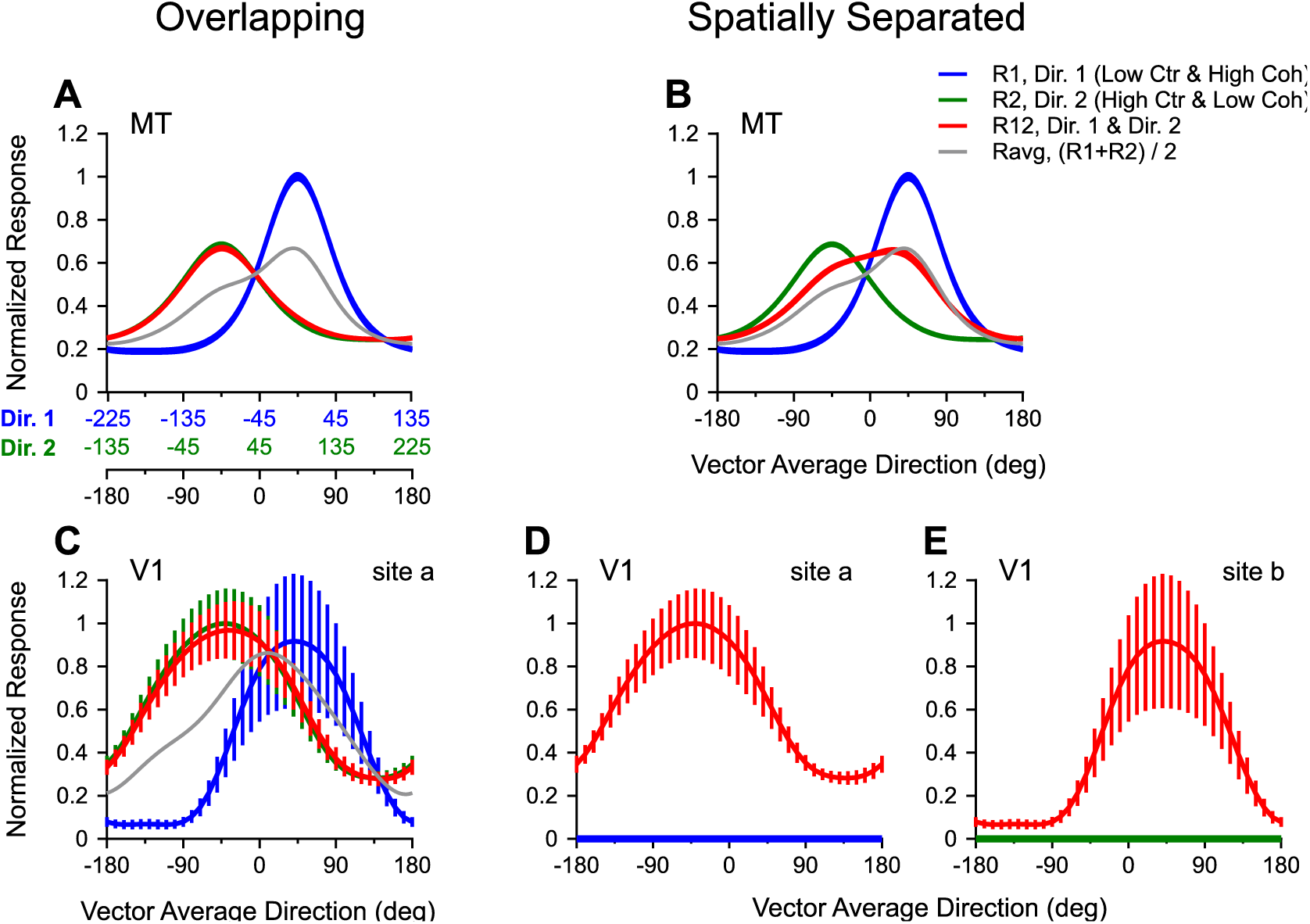
Computer simulations of direction tuning curves of MT and V1 neurons to the bi-directional stimuli used in the main physiological experiment. The visual stimuli are either overlapping (**A, C**) or spatially separated (**B, D, E**) within the RFs of model MT neurons. The two stimulus components compete in luminance contrast and motion coherence. The simulated responses to the “low contrast & high coherence” component and the “high contrast & low coherence” component are shown in blue and green, respectively. The responses to the bi-directional stimuli are shown in red. The vector average direction and the directions of individual stimulus components are labeled in the corresponding X-axes (**A**), following the same convention as in Figure 2. **A, B.** Simulated responses of model MT neurons. **C-E.** Simulated responses of model V1 complex cells. Widths of the tuning curves in A and B and the error bars in C-E represent standard deviations.

At the V1 stage of the model, the tuning curves of V1 complex cells showed a slightly greater mean peak response to the “high contrast & low coherence” component than to the “low contrast & high coherence” component (Fig. 9C). In the overlapping condition, the simulated V1 response elicited by the bi-directional stimuli was strongly biased toward the “high contrast & low coherence” component (Fig. 9C), to the extent similar to that found in model MT neuron (Fig. 9A), as measured by the weights for the component responses using the SNL model fits. The bias index (Eq. 4) for the V1 model neuron was 0.90 and that for the MT model neuron was 0.93. These simulation results suggest that the strong bias toward the “high contrast & low coherence” component found in MT is inherited from V1.

In the spatially-separated condition, the V1 response elicited by the bi-directional stimuli was the same as that elicited by the single stimulus component placed within the RFs of V1 neurons (Fig. 9D, E). Although the V1 peak response elicited by the “high contrast & low coherence” component at site *a* was slightly stronger than that elicited by the “low contrast & high coherence” component at site *b*, the MT response elicited by the bi-directional stimuli was skewed toward the “low contrast & high coherence” component, consistent with the average of the component responses (Fig. 9B). These simulation results suggest that MT response elicited by the bi-directional stimuli in the spatially-separated condition (Fig. 9B) may be due to feature competition within MT.

The response tuning curves of single MT neurons measured by varying the VA direction of the bi-directional stimuli can be mapped to the responses of a population of MT neurons that have different PDs, elicited by the bi-directional stimuli moving in a given VA direction. Figure 7 summarizes the changes of the response distributions across neuron populations at V1 and MT stages, under the overlapping and spatially-separated conditions. These results reveal the importance of neural processing at different stages of the visual hierarchy on determining how multiple visual stimuli compete within neurons’ RFs in a given brain area.

## Discussion

We have shown that how MT neurons represent multiple stimuli competing in more than one feature domain depends on the spatial arrangement of the visual stimuli. When two stimuli are overlapping, MT responses are dominated by the stimulus component that has high contrast. When two stimuli are spatially separated, the contrast dominance is abolished. Our neural data and model simulations suggest that the contrast dominance found with overlapping stimuli is due to normalization occurring at an input stage fed to MT, and MT neurons cannot overturn this contrast dominance based on their own feature selectivity. The interaction between spatially separated stimuli can largely be explained by normalization within area MT. By using multiple visual stimuli competing in more than one features domain, our study revealed how neural processing along the hierarchical visual pathway shapes neural representation of multiple visual stimuli in extrastriate cortex.

### Consideration of the effect of attention

Attention can bias neuronal responses elicited by multiple stimuli in the RF in favor of the attended stimulus (Reynolds et al., 1999; Li and Basso, 2005; Treue and Maunsell, 1996; Ferrera and Lisberger, 1997; Treue and Martinez-Trujillo, 1999; Recanzone and Wurtz, 2000; Lee and Maunsell, 2010). Although in this study the animals performed a fixation task without the need to engage goal-directed attention, could the high contrast component capture stimulus-driven attention (Corbetta and Shulman, 2002) and bias the neuronal response elicited by the overlapping stimuli? Several considerations argue against this possibility. While an abrupt stimulus onset captures attention (Yantis and Jonides, 1984), a visual stimulus that is brighter than other distractors does not automatically capture attention (Jonides and Yantis, 1988). The two stimulus components of our overlapping stimuli were turned on and started to move at the same time. The stimulus onset may automatically draw attention toward the spatial location of the overlapping stimuli, but it is unlikely to draw attention toward only the high contrast component. Furthermore, stimulus-driven attention occurs with a time delay (Nakayama and Mackeben, 1989) and its effect on neuronal responses in MT is transient, lasting for about 70 ms (Busse et al., 2008). In contrast, we found that the response bias toward the high contrast component is present in the very beginning of the neuronal responses following the onset of the static stimuli, and the bias is persistent throughout the motion period (Fig. 5). In addition, Wannig and colleagues (2007) have shown that attention directed to one of two overlapping surfaces can alter the responses of MT neurons. However, attention led to a response magnitude modulation of about 20% in MT between conditions when attention was directed to two different surfaces (Wannig et al., 2007). Even if, for some reason, the animals were consistently attending to the high contrast component throughout the stimulus presentation period in our study, the effect of attention would be insufficient to account for the nearly complete dominance by the high contrast component.

### Mechanisms underlying stimulus interactions

The primate visual system is hierarchically organized (Maunsell and van Essen, 1983; Felleman and Van Essen, 1991). The response properties of neurons in a visual area are shaped by feedforward input, as well as intra-areal and feedback processes. To understand the mechanisms underlying neural encoding of multiple stimuli, it is important to determine how these processes contribute to the RF properties in a given visual area. However, it is often difficult to disentangle the contribution of feedforward input from other neural processes. We have previously found that, in response to overlapping stimuli, MT neurons show a bias toward the stimulus component that has a higher signal strength, defined by either luminance contrast or motion coherence (Xiao et al., 2014). The response bias can be described by a model of divisive normalization. Because neurons in V1 also show a bias toward the stimulus component that has a higher contrast (Busse et al., 2009; MacEvoy et al., 2009) and divisive normalization may occur in both V1 and MT (Simoncelli and Heeger, 1998; Heuer and Britten, 2002), it was unclear how the feedforward input from V1 contributed to the response bias found in MT.

In this study, we are able to differentiate the impact of feedforward input from other neural processes on the response properties of MT neurons. Our results suggest that neurons in V1 may respond more strongly to the “high contrast & low coherence” component than to the “low contrast & high coherence” component used in our experiment, due to V1 neurons’ sensitivities to contrast and coherence. When two stimuli overlap, the responses of V1 neurons elicited by both stimulus components may already show a strong bias toward the “high contrast & low coherence” component due to divisive normalization in V1 (Fig. 9C). MT neurons are no longer able to remix the stimulus components according to their own sensitivities to contrast and coherence. In other words, MT neurons inherit the response bias toward the high contrast component from their input. When two visual stimuli are spatially separated, MT neurons receive inputs from two different pools of V1 neurons and each neuron pool responds to only one stimulus component (Fig. 7B). The neuronal responses elicited by the two stimulus components remain separated in V1. MT neurons can mix the responses elicited by the two stimulus components via spatial and directional pooling and divisive normalization within MT. As a result, the mixing in MT may well reflect the sensitivities of MT neurons to different stimulus features. Our model simulations make predictions regarding how V1 neurons respond to multiple competing stimuli (e.g. as shown in Fig. 9C), which can be tested in future physiological study.

### Implications on normalization and encoding of multiple visual stimuli

Our finding that the response weighting for competing stimuli depends on the spatial arrangement provides a new perspective on the well-established normalization model (Carandini and Heeger, 2011). The basic form of normalization equations (Eqs. 5-6) predicts that the response weight for a stimulus component increases with its signal strength, but does not consider the spatial arrangement of the visual stimuli. We made a surprising finding that MT response to overlapping stimuli cannot be predicted by the population neural responses in MT elicited by the individual stimulus components. One must consider the neural computations occurring along the hierarchical visual pathway.

Majaj, Carandini, and Movshon (2007) showed that pattern-direction selective neurons in MT characterized by overlapping drifting gratings (i.e. plaid) do not integrate the directions of the component gratings when they were spatially separated within the RF, suggesting that the computation underlying pattern-direction selectivity in MT is local. Different from the plaid, the overlapping random-dot stimuli used in our study elicit the percept of motion transparency. We showed that changing the spatial arrangement of visual stimuli can have a substantial impact not only on motion integration but also on the competition between multiple stimuli. Our results revealed that contrast has a dominant effect in determining stimulus competition within a local spatial region when multiple stimuli differ in more than one feature domain. When visual stimuli are spatially separated, the effect of contrast is substantially reduced.

A seminal model involving MT neurons pooling inputs from V1 and divisive normalization in both V1 and MT has been successful in explaining a range of experimental results of MT responses (Simoncelli and Heeger, 1998; Rust et al., 2006). However, the model in its original form does not specify how features are spatially integrated and it does not differentiate overlapping and spatially separated stimuli (Majaj et al., 2007). In our study, we adapted this model to simulate both overlapping and spatially separated conditions and showed that the framework can explain our main physiological findings. Also using this model, Busse, Wade, and Carandini (2009) previously demonstrated the impact of response normalization in V1 on neural response in MT. They showed that, by making the contrasts of two drifting gratings of a plaid to be unequal, the response of a model MT neuron changed from representing the pattern motion of the plaid to mostly representing the higher-contrast grating component, likely due to contrast normalization in V1 (Busse et al., 2009). However, the MT response elicited by the higher-contrast grating alone could also be greater than that elicited by the lower-contrast grating. The model-predicted response bias toward the higher-contrast component in MT may also be contributed by response normalization within MT, akin to our experimental result obtained using random-dot stimuli with unequal contrasts (Xiao et al., 2014). In comparison, our current study provides unequivocal new evidence on how responses in MT are shaped by the hierarchical network. By using two stimuli competing in more than one feature domain, we demonstrated neurophysiologically and computationally the substantial impact of stimulus competition in the input stage on the neuronal responses in MT and how that impact changes with the spatial arrangement of visual stimuli. Our finding may also apply to other visual areas in the hierarchical network, including those in the ventral visual stream where response normalization has been well documented.

## Acknowledgment

We thank Jianbo Xiao for assistance on electrophysiological recordings and data analysis and Bryce Arseneau for technical support. This work was supported by National Institutes of Health Grant R01EY022443, and in part by the Office of the Director, National Institutes of Health Grant P51OD011106 to the Wisconsin National Primate Research Center, University of Wisconsin-Madison. This research was conducted at a facility constructed with support from Research Facilities Improvement Program grant numbers RR15459-01 and RR020141-01.

